# Cryo-Electron Tomography 3D Structure and Nanoscale Model of *Arabidopsis thaliana* Cell Wall

**DOI:** 10.1101/492140

**Authors:** Purbasha Sarkar, Michal Kowalczyk, Salil Apte, Edgar G. Yap, Jyotirmoy Das, Paul D. Adams, Chandrajit Bajaj, Pablo Guindos, Manfred Auer

## Abstract

Using cryo-electron tomography of vitrified sections of one month-old *Arabidopsis thaliana* inflorescence stem tissue, we visualized primary and secondary cell walls of xylem tissue. Extensive quantitative and statistical analysis of segmented 3D tomographic data allowed geometrically idealized 3D-CAD model building of prototypic microfibrils, cross-links, and their supramolecular microfibril 3D organization. We propose a prototypic microfibril model where a cellulose core is heavily decorated by a thin sheath of hemicellulose with infrequent but sturdy hemicellulose-based cross-links. Such prototypic microfibrils then adopt a rather unexpected 3D supramolecular organization of high order and complexity. We discuss a possible new role for lignin in plant cell walls at low concentrations with lignin not acting as a matrix but rather as a reinforcement of microfibrils and cross-links. Extensive computational simulations of mechanical properties further revealed that this 3D organization of the cell wall is not optimized for load bearing but instead for flexibility and ductility.

**One Sentence Summary:** Cryo-electron tomography and mechanical simulations revealed cell wall 3D architecture, optimized for flexibility/ductility.

---

Cell walls provide mechanical strength and protection to plants, while being flexible to support their growth and development and have been studied for over eight decades (1-6). The current model of plant cell walls portrays a framework of ~3.5 nm thick cellulose microfibrils, interconnected by single-strand hemicellulose thus forming a rather loose 3D “long-tether” network, with hemicellulose adhering to significant stretches of the microfibrils (7-14). Other prominent cell wall components, pectins are predominantly found in the middle lamella (ML) adhering two adjacent cells (15), whereas phenolic lignin polymers are thought to form a hydrophobic matrix in between the cellulose-hemicellulose network (15, 16). The most commonly discussed cell wall model has microfibrils alternating between two sharply distinct orientations, not unlike textile fabric (3, 7, 11), however alternative models have been proposed (1-2, 17-19).

Numerical simulation of mechanical properties using approximation techniques such as the finite element method (20) require realistic 3D cell wall structures and models, and to date have been based on the assumption of pseudo-random fiber networks (21-23). Model building allows virtual testing of the different cell wall components and alternative 3D configurations. As for fiber-reinforced composites, homogenization techniques can be combined with finite element methods to extract global properties such as stiffness of the wall.

Here we used cryo-electron tomography vitreous sections of 1-month old *Arabidopsis thaliana* inflorescence stems (24) to reveal the 3D nano-architecture of microfibrils and their cross-connectors, as well as the supramolecular 3D organization of primary and secondary walls. Extensive statistical volumetric analysis and model building, followed by mechanical properties simulations suggest that the secondary cell walls we examined are not optimized for maximal load bearing but instead for flexibility and ductility with superior load transfer capabilities as well as elastic and viscoelastic energetic capacities.

## Cell Wall Building Blocks

1-month old *Arabidopsis thaliana* inflorescence stems (Fig. 1A-B) have different cell wall types (Fig. 1C), including thin xylem parenchyma (XP) cell walls with only primary walls and thick xylem tracheary (XTE) elements cell walls with both primary and secondary walls. Figure 1D and 1E show single ~1nm thin slices through 3D reconstruction of vitreous sections of XP and XTE cell walls cut from self-pressurized, ultra-rapidly frozen young stem segments (24), with corresponding 3D renderings shown in Fig. 1F and 1G, respectively.

**Fig 1.**
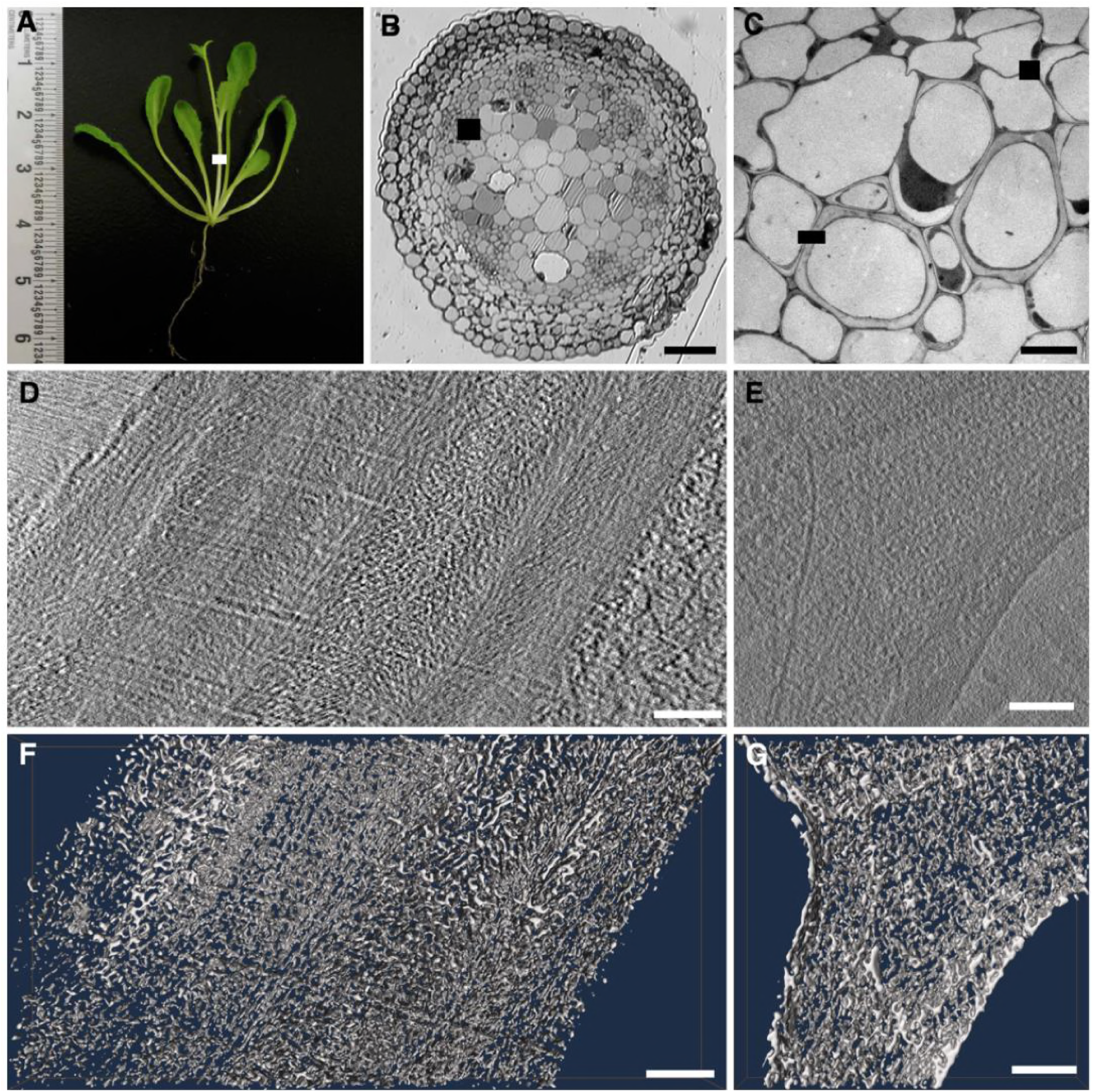
Cryo-electron tomography of thick xylem tracheary elements cell walls and thin xylem parenchyma. A) 1 month-old *Arabidopsis thaliana* inflorescence stem tissue. The white box marks a stem segment that was taken for cryo-electron tomography. B) Cross-section through the stem showing the different tissue and cell types in the stem. The black box marks a tissue region similar to the one used for cryo-EM imaging. Note that the section displayed is a Toluene Blue-stained semithin section cut from a high-pressure frozen, freeze-substituted and resin-embedded samples imaged by optical light microscopy in order to provide an overview of the tissues and cells in the plant stem organ. Scale bar = 50 μm. C) Electron micrograph of a ~25 μm by ~25 μm tissue region of interest showing both a thick xylem tracheary elements cell wall (left box) and a thin xylem parenchyma cell wall (right box). Scale bar = 5 μm. D) Zero degree projection view cryo-electron microscopy of thick xylem tracheary elements cell wall, containing primary and secondary cell wall. E) Zero-degree projection view cryo-electron microscopy of thin xylem parenchyma cell wall, containing only primary cell wall. F) 3D rendering of a ~50 nm thin slab of a cryo-tomogram of xylem tracheary elements cell wall. G) 3D rendering of a ~50 nm thin slab of a cryo-tomogram of xylem parenchyma cell wall. Scale bars = 100 nm.

Measuring the diameter of individual microfibrils (Fig. 2A, n=150 for each cell wall type), we observed peaks at 3.5 ± 0.5 nm and at 5 ± 0.5 nm (Fig. 2B), which in accordance with the current microfibril model, may correspond to a round cellulose core that in many places is surrounded by a very thin sheath of hemicellulose. Some microfibrils appear elliptical with up to ~9 nm thicker portions (Fig. 2B-C) that may be assigned to additional hemicellulose, pectin, or lignin. This assignment was further supported by the experimental removal of pectins and hemicelluloses (24-25) from primary cell walls (Ext. PCW) by treatments with 0.5% ammonium oxalate and 4% NaOH prior to dehydration and resin-embedding (Fig. 2B). Cellulose, hemicellulose and pectin cannot be directly distinguished in the density maps due to their chemical similarity. We will use the term matrix polychaccarides instead of hemicellulose, since pectin’s contribution to microfibrils and cross-links cannot be ruled out, despite the overall low pectin concentration and expected localization of pectin to the middle lamella.

**Fig 2.**
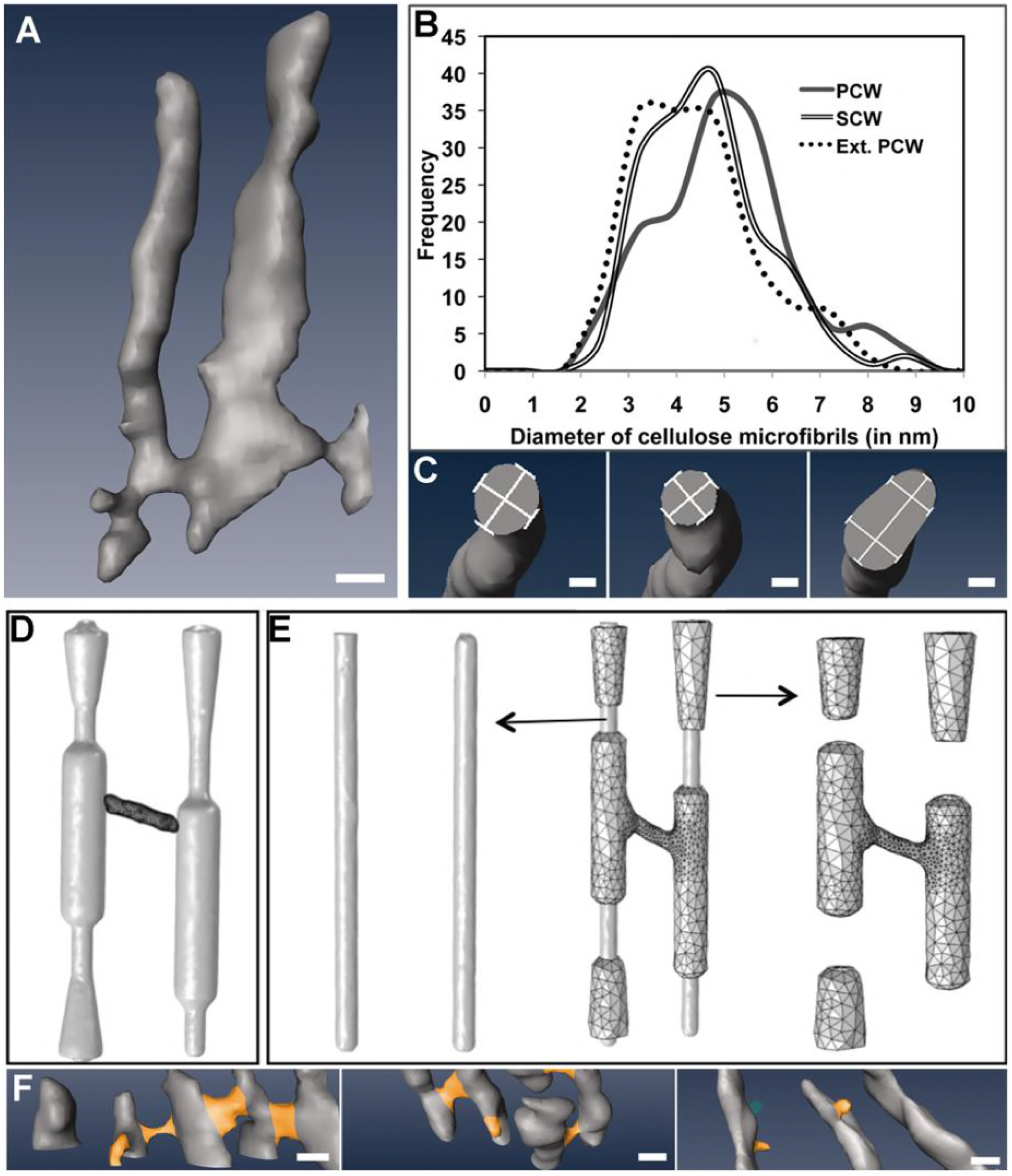
Statistical analysis based 3D modeling of microfibrils and cross-links A) 3D rendering of two neighboring microfibrils in the cryo-tomogram. Scale bar = 5 nm. B) Plot of the statistical analysis of the frequency of microfibril diameter reveals a peak at 3.5 nm and 5-5.5 nm, and a small shoulder to dimensions up to 9 nm. C) Gallery of 3 different microfibrils cross-sections illustrating the numbers obtained in Fig 4B) with microfibrils being round at ~3.5 nm (middle panel), oval shaped with a diameter of ~3.5 nm by ~5-5.5 nm (left panel), reaching up to 9-10 nm (right panel). Scale bars = 2 nm. D) Hypothetical 3D-CAD model with all of cellulose residing in the microfibrils and hemicellulose confined to cross-links only. E) Alternative 3D model of two adjacent microfibrils (middle) that contain a cellulose core (left), surrounded by a hemicellulose sheath as well as a cross-connection between adjacent microfibrils (right). F) Occurrence of cross-links in PCW (left), SCW (middle) and chemically extracted PCW (right). Scale bars = 5 nm.

Statistical analysis of microfibrils and cross-connector thickness allowed a comparison between the volumes occupied by cellulose and matrix polysaccharides, respectively, and to build idealized models: one where microfibrils are exclusively made of cellulose, with all matrix polysaccharides residing in the cross-links only (Fig. 2D) and another scenario where cellulose strands form a 3.5 nm microfibril core, surrounded by a thin matrix polysaccharides-based sheath and matrix polysaccharides form sturdy cross-links (Fig. 2E). The matrix polysaccharides-to-cellulose volume ratio of the first scenario is ~0.07 and 1.3 for the second scenario, which is in good agreement with bulk analysis estimates from *Arabidospsis thaliana* root and leaves (26). While we realize the limitations of comparison with bulk mass ratio biochemical analysis, our core-and-sheath model seems more likely, and is agreement with previous findings that significant stretches of matrix polysaccharide strands are closely aligned with the elementary fibril cellulose core (9, 10, 12, 27-28). This arrangement would allow hemicellulose sheaths and cross-links to slide along the cellulose core under shear force, whereas rigid, highly localized connections would likely break and has significant implications for force transmission between adjacent microfibril layers as will be discussed below. Bridge-like cross-linkers (n=100 for each type of cell wall) appeared short (typically 4-6 nm) and thick, indicating bundling of multiple matrix polysaccharide strands. Within the z=50 nm section-height examined in cryo-tomograms, we found on average for 4 and 3 out of every 5 microfibrils for PCWs and SCWs, respectively. Cross-links were absent in chemically extracted PCWs (Fig. 2F).

## Supramolecular 3D Cell Wall Organization

As shown in Figure 3A-B xylem tracheary element (XTE) cells feature multiple layers, with the middle lamella (M) being sandwiched between primary cell wall layers (P), which are flanked by three secondary cell wall segments (S1, S2, S3) on one side and one (S) secondary cell wall segment on the other side, with intermittent transition zones (T). Xylem parenchyma (XP) cells lack secondary cell walls layers (Fig 3C-D). To determine whether texture differences across the cell wall visible in cryo-EM projection images of XTE cell walls (Fig 3A) could be attributed to differences in microfibril 3D orientations, we measured the average tilt angle of microfibrils in consecutive radial microfibril layers. We found in each of the S1, S2 and S3 regions of SCWs ~15 consecutive parallel microfibril layers (Fig. 3B) that were off-set from axial orientation by either plus or minus ~27° (with slight variations being present in the tilt angle value for each microfibril). Between each of the S regions and between the S and P region, we found a three-layer transition zone (T), where microfibrils were oriented axially. Likewise, in primary cell walls in both XTE and XP cells (Fig. 3C-D) most microfibrils were oriented axially (parallel to the growth axis).

**Fig 3.**
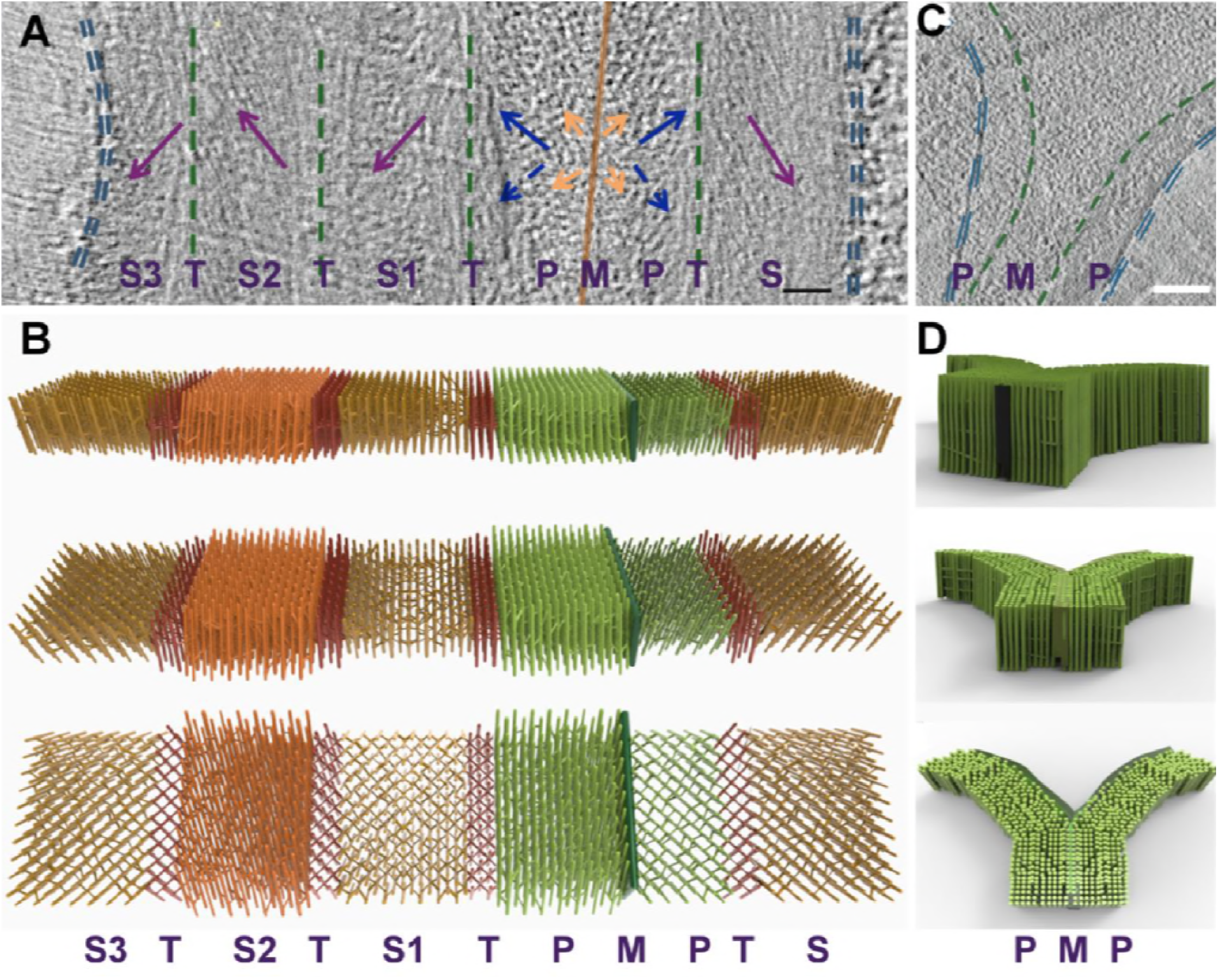
Supramolecular 3D organization of microfibrils in xylem tracheary elements and xylem parenchyma cell walls. A) Zero-degree projection image of xylem tracheary elements cell wall, showing two adjacent cells with both secondary and primary cell walls. Note that one of the two adjacent cells has several secondary cell wall subregions, called S1, S2 and S3 and transition zones (T). On the other side of the middle lamella (M) is the primary cell wall P, a transition zone T as well as one secondary cell wall region.Scale bar = 100 nm. B) Idealized model of a xylem tracheary elements cell wall at side view (top), slanted 45° view (middle) and an en-face view of microfibrils (bottom). Note that microfibril orientation differs in S1, S2 and S3, with 15 layers of microfibril featuring an average angle of minus ~27°, plus ~27° and minus ~27° off the longitudinal axis (plant elongation direction). S3, S2, S1 and the Primary Cell Wall (PCW) regions are separated by a three layer-transition zone with axial microfibril orientation. The pectin-rich middle lamella is depicted in the idealized model as a single plane separating the cell walls of two adjacent cells. C) Zero-degree projection image of xylem parenchyma cell walls, showing two adjacent cells with only primary cell walls. Scale bar = 100 nm. D) Idealized model of a xylem parenchyma cell wall at side view (top), D2) slanted 45° view (middle) and an en-face view (bottom). Note that in PCW microfibril orientation is mostly axial.

## Mechanical Cell Wall Properties

To determine what effect the supramolecular 3D organization had on the mechanical properties of primary and secondary cell walls, we resorted to computational simulations (Fig. 4). First we examined the mechanical properties of the rather complex tomography-derived 3D volume of the complex XTE walls (Fig. 4A) and of simplified and idealized 3D-CAD models (Fig. 4B), which allowed for the calculation of axial load and shear forces for different wall models. We estimated the overall mechanical stiffness on both the experimentally determined volumes (Fig. 4C) and CAD-model idealized models (Fig. 4D), using homogenization, as is a routine approach in composite and porous material research.

**Fig 4.**
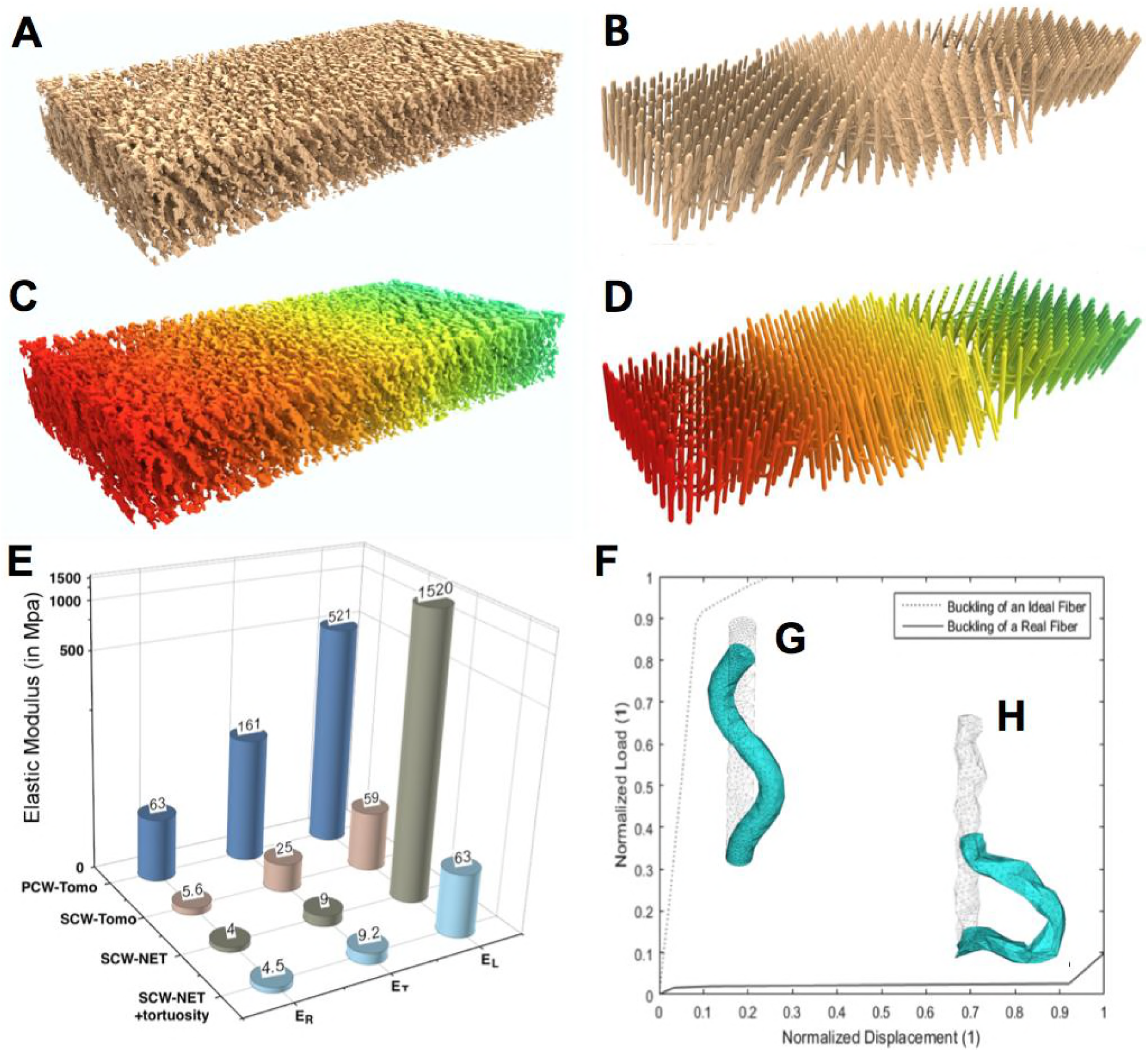
Computational analysis of cell wall mechanical properties. (A, B) Supramolecular structure segmented from the tomographic reconstruction (A) and a geometrically idealized cell wall model (B) both including only the S1-T-S2-T-S3 portions of the idealized model. (C, D) Deformation of the cell wall under radial pressure, for both the segmented volume (C) as well as an idealized model (D), revealing a linear distribution of force throughout the secondary cell wall. (E) Graphic depiction of the Young elastic moduli (radial, longitudinal and tangential) for primary cell walls, secondary cell wall regions segmented from the tomographic 3D data as well as idealized models of the secondary cell walls, before and after introduction of tortuosity. (F, G, H) Load bearing behavior of microfibrils under buckling due to axial stress, revealing significant decrease of failure load but very high increase of ductility (F), when moving from an idealized straight microfibril (G) to a tortuous fiber (H).

### Mechanical properties of tomographic cell wall density maps

We assumed a 300×100×50nm^3^ tomographic representative volume element (RVE) of SCW to be made of orthotropic material (Supplementary text; Tables S1 and S2), and considered Young’s modulus of 30 and 10 GPa for cellulose and hemicellulose, respectively (21). We found the effective stiffness of the SCW tomographic volume (SCW-Tomo) to be E_R_=5.6 MPa, E_T_=25 MPa and E_L_=59 MPa, along the principal material axes (radial, transverse, longitudinal) respectively (Fig. 4E). Likewise, we also performed a mechanical analysis of the PCW tomographic volume (PCW-Tomo), and found the stiffness to be E_R_=63 MPa, E_T_=161MPa and E_L_=521MPa (Fig 4E). We attribute the higher stiffness of the PCW-Tomo compared to the SCW-Tomo to a smaller void fraction, which was surprisingly large (PCW: ~72%; SCW: 80%). The void volume is the space not taken up by microfibrils or cross-links, which is likely filled with water and soluble small molecules. Our cryo-tomographic analysis suggested the absence of an extensive lignin network matrix, as oligomers or polymers larger than 3 nm should be visible in our cryo-tomograms. We modeled lignin as ball-like objects ranging from 0.5-3 nm to be added into the SCW-Tomo volume at various concentrations from 0-30% of the total volume fraction (Fig. S2 A) and found that at low lignin concentrations there wasn’t enough material to form a continuous matrix, instead lignin may reinforce the cellulose-hemicellulose macromolecular network, as even small increases in lignin concentrations has significant effects on the Young moduli, which showed quadratic increases in all directions (Fig. S2G). We submit that the role of lignin at low concentrations is not to act as a matrix but to reinforce the microfibril-marix polymer framework, thus enhancing cell wall’s overall stiffness.

### Comparison idealized 3D-CAD model with tomographic density maps

We considered four idealized modeling approaches (see Supplemental material and Fig. S1) to both primary and secondary cell wall, and chose the NET model for further simulations. The calculated stiffness in SCW-NET (E_R_=4MPa, E_T_=9 MPa, E_L_=1520 MPa) was in good agreement with the homogenized SCW-Tomo results, except for the stiffness in the axial direction, which was ~25 fold higher (Fig. 4E). This discrepancy led us to an in-depth examination of microfibril shapes. We realized that microfibrils were not straight rods but had a wavy appearance, which corresponds to a weakening of the microfibril in the axial direction. To model such imperfections we introduced a tortuosity (twist) in our idealized microfibril model, which had a very small influence in the transverse direction (E_R_=4.5 MPa, E_T_=9.2 MPa), but resulted in stiffness drop in the axial direction (E_L_=63 MPa), bringing it in close agreement with the SCW-Tomo results in all three directions (Fig. 4E).

Nonlinear buckling and viscoelastic analyses of the fibers with and without tortuosity (Fig. 4F) revealed that that tortuous fibers withstand ~50 times lower buckling load compared to straight fibers (Fig. 4G-H) but can undergo ~9-times larger deformations before collapse and store up to 43% more elastic energy as they deform. We hence, conclude that imperfections in microfibril structure, while significantly reducing the stiffness of the cell wall, increases its ductility as well as the dissipation of elastic and viscoelastic energies, which could be crucial for plants to prevent breakage against extreme loading (such as in high winds).

### Supramolecular 3D organization of microfibrils across the cell wall

We further examined the effect of the alternation of microfibril orientation (plus 27° or minus 27°) for each of the three 15-microfibril layers in the S1, S2 and S3 SCW and the 3-layer transition zone in axial microfibril orientation (Fig. S3A). The minus/plus/minus 27° deviation from an axial orientation caused a dramatic stiffness drop in the axial direction, but also a 200-fold increase in the elastic strain energy density. Without transition zones (18 layers that are minus/plus/minus 27° inclined, Fig. S3B), axial stiffness was further reduced by a factor of 4, with an increased elastic energy storage of 30%, whereas if all microfibrils (54 layers) were inclined by 27° (Fig. S3C), axial stiffness decreased 27-fold and elastic energy storage increased by 800%. The observed minus/plus/minus configuration is a quasi-symmetric and balanced composite (each ply has an opposite-aligned counterpart), thus decoupling membrane (in-plane) and bending (out-of-plane) cell wall mechanical responses (29). This could be vital for plants to prevent cell deformation during growth and external loading. Spiral spring-like configurations, e.g. in cylindrical spiral reinforced concrete columns (30), are found in civil, mechanic, automotive and aerospace engineering for their superior ductility and outstanding energy absorption capacity of impact loads, mainly due to multiple failure mechanisms (31).

## Acknowledgement

We thank Drs. Carragher, Potter, Quispe, Jacovetty, Cheng from National Resource for Automated Molecular Microscopy (NRAMM), Csencsits, Bosneaga, Downing (LBNL), McDonald, Zalpuri (UC Berkeley). We acknowledge support by Energy Biosciences Institute (grant 007G18). Work was continued at Joint BioEnergy Institute, supported by the U.S. Department of Energy, Office of Science, Office of Biological and Environmental Research, through contract DE-AC02-05CH11231 between Lawrence Berkeley National Laboratory and the U.S. Department of Energy. The United States Government retains and the publisher, by accepting the article for publication, acknowledges that the United States Government retains a non-exclusive, paid-up, irrevocable, worldwide license to publish or reproduce the published form of this manuscript, or allow others to do so, for United States Government purposes. MA acknowledges support of the cryo-electron microscopy at LBNL by the NIH/GMS grant P01GM051487.

The authors declare no financial and non-financial competing interests.

## Supplementary Materials

**Fig S1.**
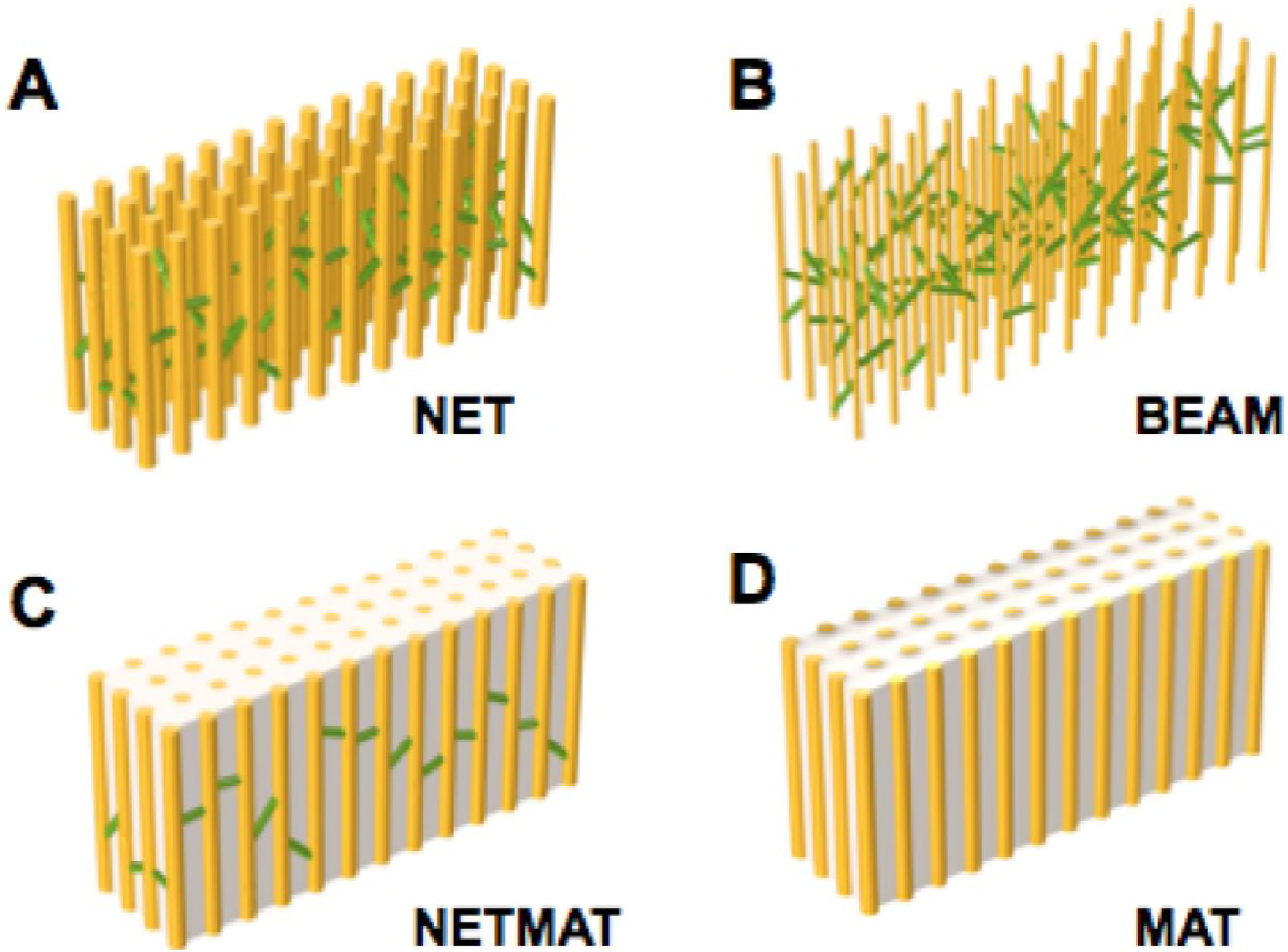
Choice of Modeling Approach. We considered four idealized modeling approaches: (A) NET is a network model that fully accounts for the 3D volume of both microfibrils and cross-links B) BEAM is a simplified approach based on Timoshenko beam mechanical theory (C) NETMAT is similar to NET, with an added a matrix in between the fibers and (D) MAT, which is similar to NETMAT but does not contain inter-microfibrillar cross-connectors.

**Fig S2.**
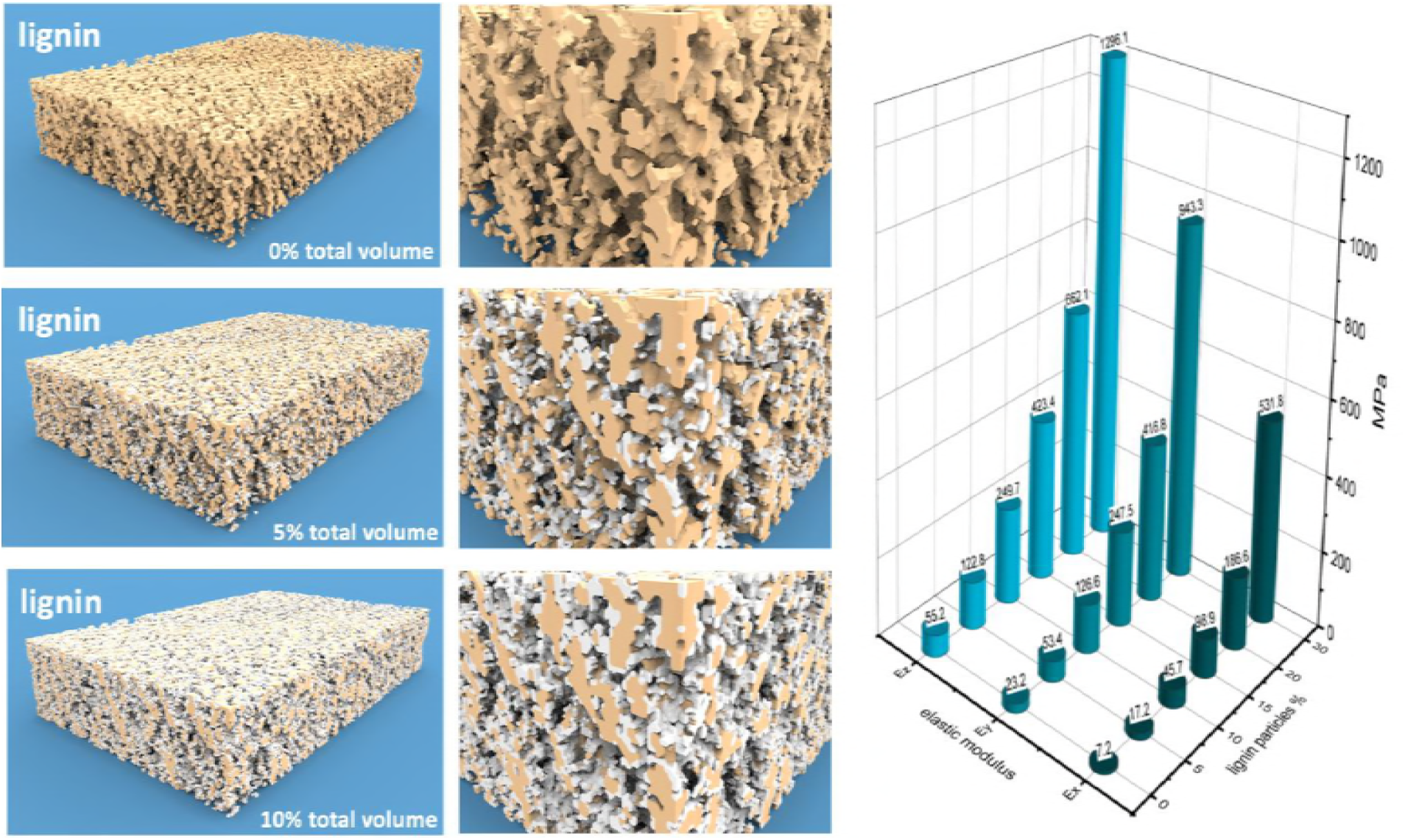
Modeling of lignin distribution in secondary cell wall (A, D) Secondary cell wall with an assumed 0% lignin. 20% of the total volume available is occupied by the cellulose/hemicellulose network. (A) overview (D) close-up detail. (B, E) 5% total volume of lignin (equivalent to a 15-20% dry weight) is added to the segmented map displayed in A. (B) overview and (E) close-up detail reveal vast amount of space remains unoccupied, inconsistent with a dense matrix in which microfibrils are embedded and thus mechanically connected. Increasing the lignin to 10% of total volume (resulting in a ~30-35% dry weight) leads to much more densely connected network, which is consistent with a matrix at these much higher lignin content. (C) overview (F) close-up detail. Note that the corresponding dry-weight is an estimate making assumptions on similarity in density of lignin and cellulose/hemicellulose matrix. (G) Comparison of elastic moduli for radial (Er), tangential (Et) and longitudinal (EL) force loading.

**Fig S3.**
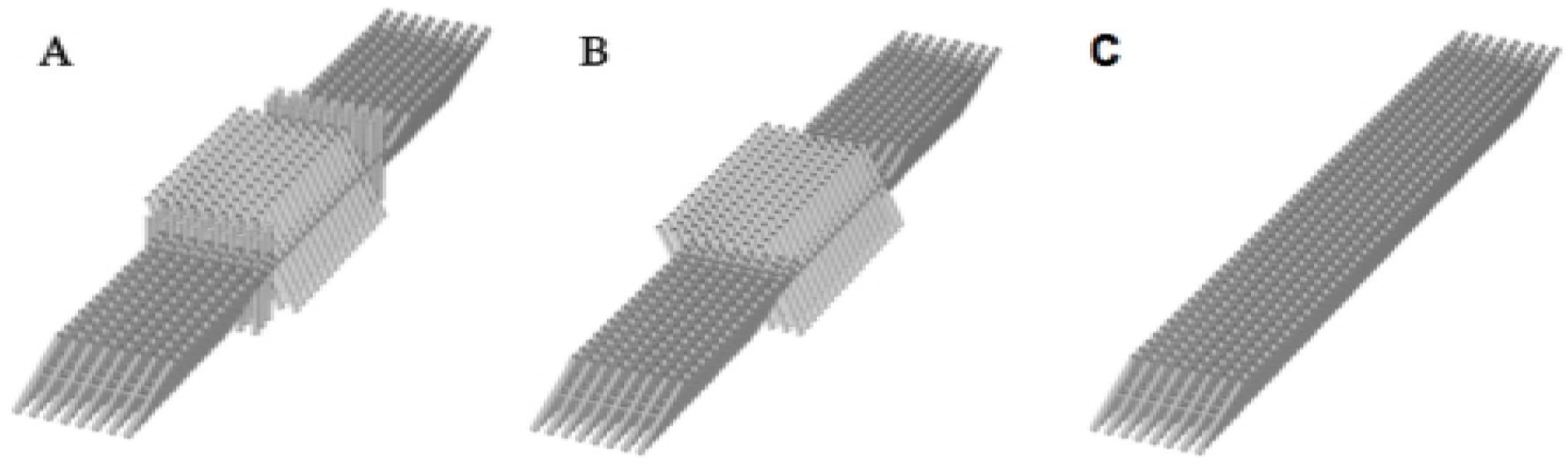
Hypothetical XTE cell wall models with varying supramolecular organization. A) Cell wall model with 15 microfibril layers each in the S1, S2 and S3 SCW that were inclined minus 27°, plus 27° and minus 27°, respectively, and that were separated from each other by a 3 microfibril layer transition zone with axial microfibril orientation. B) Cell wall model without transition zones, and 18 layers of microfibrils that are minus/plus/minus 27° inclined. C) All microfibrils (54 layers) are all inclined by 27°.

**Table S1.** Differences on the calculated properties calculated by homogenization method on differently sized density map fragments.

| Size of the Density Map Fragment Used on the Calculation ($\text{[nm}^3\text{]}$) | | Absolute Difference Between RVE A and RVE B on the Calculated Mechanical Properties (%) |
| --- | --- | --- |
| RVE A | RVE B |  |
| 200x200x50 | 300x100x50 | 40 |
| 300x100x50 | 400x350x50 | 15 |
| 300x100x50 | 300x300x100* | 13 |
| 300x100x50 | 300x300x200* | 12 |

**Table S2.** Average strain equivalence errors of the different elastic laws used in homogenization.

| Elastic Constitutive Law Used in Homogenization | Average Strain Equivalence Error |
| --- | --- |
| Isotropy | 67% |
| Transverse isotropy | 20% |
| Orthotropy | 6% |

## SUPPLEMENTAL MATERIAL

### Materials and Methods

#### Plant Material

Wild type *Arabidopsis thaliana* (Arabidopsis) seeds from the Colombia ecotype (Col 0) were sterilized in 30% bleach, 0.02% Triton and vernalized at 4 °C in water for 48 hours. They were germinated on 0.7% agar plates containing 0.5x Murashige and Skoog medium for 10 d at 21 °C under continuous light in a growth chamber. The seedlings were then transferred to pots containing soil mixture and placed in a growth chamber programmed for a 16 h light/8 h dark cycle at 21 °C. Stem tissue from ~1 month old plants that had newly growing inflorescence stems (2-3 cm long) were used for electron tomography. Stem segments from freshly collected samples were used for electron tomography.

#### Cryo-electron tomography of *Arabidopsis thaliana*

~2 mm long stem segments of *Arabidopsis thaliana* (Col 0) were frozen in sealed copper capillary tubes in 20% dextrane by self-pressurized rapid freezing method (32) and sectioned at −160°C using a Leica EMUC7 ultramicrotome with a Leica EM FC7 cryo chamber attachment (Leica Microsystems Inc.). Ribbons of nominal ~90 nm ultrathin cryo-sections were manipulated by hand with an eye lash and placed onto carbon-coated lacey formvar grids and attached to the grids using a Leica EM CRION ionizer. Single-axis cryo tilt series were collected from −60° to +60° with 2° increments under low dose conditions on two different set-up. Some datasets were collected on a Tecnai F20 TEM (Thermofisher, Inc, Hillsboro, OR, USA) with a 4K × 4K Gatan Ultrscan 4000 CCD camera (Gatan Inc., Pleasanton, CA, USA) and Leginon (33) at 120 kV and a voxel size of 0.44 nm. Other datasets were collected on a JEOL JEM–3100FFC TEM (JEOL Ltd, Akishima, Tokyo, Japan) equipped with a field emission gun electron source operating at 300kV, an in-column Omega energy filter (JEOL), a cryo-transfer stage and a Gatan 795 2K×2K CCD camera and SerialEM (34) and a voxel size of 1.1 nm voxel size. Images were aligned by patch-tracking method in IMOD. Reconstruction of all tomograms was done with IMOD (The Boulder Laboratory for 3D Electron Microscopy of Cells, University of Colorado Boulder, CO) using the back-projection method (35-37) Cryo-tomograms collected on a 4K × 4K camera were binned by 2 to obtain a voxel size of 0.867 nm, to be comparable with the other tomograms. All tomograms were subjected to image filtering (nonlinear anisotropic diffusion filter) to improve contrast.

#### Pectin and hemicellulose removal from *Arabidopsis thaliana*

~2 mm long stem segments were fixed in 4% paraformaldehyde, 2% glutaraldehyde in 0.03 M phosphate buffer (pH 7.4) containing 0.5mg/ml ruthenium red overnight at 4 °C. Samples were then rinsed in the same buffer and consecutive stem segments from the same plants were treated in parallel with three different treatments: (1) Control - no chemical treatment; (2) 0.5 % ammonium oxalate at 60° C for 48 hours for pectin removal; and (3) pectin removal as (2) followed by 4 % NaOH at RT for 96 h to remove the majority of hemicelluloses and any non-cellulosic polysaccharides. All samples were rinsed in distilled water and then fixed in 0.1% osmium tetroxide with 0.5mg/ml ruthenium red for 1 h at RT. The samples were then dehydrated in acetone series and infiltrated in Epon-Araldite resin-acetone series using Leica EM AMW automatic microwave tissue processor. Samples were incubated overnight in 100% resin and then polymerized at 60°C for 2-3 days.

#### Ultrathin sectioning and electron tomography of resin embedded samples

150 nm thick sections were cut from resin embedded samples using Leica UC6 ultramicrotome (Leica Microsystems Inc.). All sections were labeled with 5 nm gold fiducials for 4 mins each on both sides followed by several washes in distilled water. The sections were then post stained with 2% uranyl acetate in methanol for 5 mins, followed by Reynold’s lead citrate solution for 2 mins. After locating areas of interest, dual axis tilt series were collected from +65° to −65° with 1° increments on a Philips Tecnai F12 TEM (FEI) with 2K × 2K Gatan Ultrascan 1000 CCD camera and SerialEM software (36, 37), at 120 kV accelerating voltage and a voxel size of ~0.8 nm. Images were aligned by tracking the fiducial markers in IMOD. Reconstructions of tomograms were done using the weighted back-projection method in IMOD.

#### Segmentation and image analysis

Segmentation of all tomograms was done by the ‘threshold segmentation’ method in Amira (Thermofisher Inc, Hillsboro, OR, USA). A triangular mesh surface was generated in Amira with the ‘unconstrained smoothing’ option, for visualization and quantitative geometric analysis of the cell wall components. Dimensions of long filamentous structures, the distance (center-to-center) and the shortest gap between the filaments, angle of microfibrils, and the dimensions of short bridge-like cross-links joining the long filaments were measured for each tomogram volume in Amira.

#### Simulation

Geometry of the ideal models was mostly generated with Matlab (MathWorks, Inc, Natick, MA, USA), mechanical analysis of the numerical models was mainly performed with COMSOL (COMSOL AB, Stockholm, Sweden) and most postprocessing was set up with Python scripts in Paraview (Sandia National Laboratory and Kitware Inc, Los Alamos National Laboratory, Los Alamos, NM, USA). Some exceptions were the nonlinear buckling analysis which was performed in ANSYS (Swanson Analysis Systems Inc., Canonsburg, PA, USA), the homogenization of the cell wall and the effect of lignification which was carried out with the Software Geodict (Math2Market GmbH, Kaiserslautern, Germany) as well as the masking of the wall to differentiate hemicellulose out of cellulose, which was conducted with VOXELCON (Quint Corporation, Tokyo, Japan).

##### Mechanical properties of experimental cell wall density maps

We applied homogenization, based on the strain equivalence principle, at different fractions of the measured cell wall, with representative volume elements (RVE) being loaded in different directions (radial, axial, shear forces). Assuming a homogeneous material this test results in predictions of the mechanical properties of entire cell walls. We used a 300×100×50 nm RVE, as larger RVEs gave very similar results, whereas smaller RVEs gave substantially different results (Table S1). The suitability of distinct elastic constitutive laws used in the homogenization calculations showed that that average strain equivalence error is low (6%) only in orthotropy, and hence is ideal for modeling the cell walls (Table S2).

##### Ideal models and comparison with the experimental density maps

For our simulations we considered four idealized modeling approaches (Fig. S1): 1) a network model that fully accounts for the 3D volume of both microfibrils and cross-links (NET); 2) a simplified version of NET (BEAM), using Timoshenko beam mechanical theory (38-39); 3) a model, with an added a matrix in between the fibers, we call NETMAT; and 4) a model similar to NETMAT but without intermicrofibril cross-links, we call MAT. We assumed the same inputs for all four models, and varied the elastic moduli of the interfibrillar matrix from 0.1 to 10 GPa. We calculated the elastic (Young) moduli (radial, longitudinal, tangential and shear), which provides an estimate of the stiffness of the cell wall, as well as the elastic strain energy density stored in the cellulose/hemicellulose and matrix, and compared with the results from the experimental map. Despite having been used frequently in previous computational simulations we found that BEAM vastly underestimates the stiffness mostly because cross-links lengths in this model are highly exaggerated and torsion is not accurately modelled. For NETMAT and MAT, the stiffness increases by up to two orders of magnitude in the transverse direction, with most of the elastic energy being stored in the matrix rather than the fibers, rendering the 3D organization of microfibrils and cross-links nearly meaningless. We chose the NET model for our idealized model simulations. The calculated stiffness in NET (E_R_=4MPa, E_T_=9 MPa, E_L_=1520 MPa) were in good agreement with the homogenized tomography model results, except that the stiffness in the longitudinal (axial) direction was ~25 fold higher (Fig. 4E). However, if tortuosity is considered in the NET model (see below), it has a very limited influence in the radial and transverse directions, but the stiffness drops drastically in the axial direction (E_R_=4.5 MPa, E_T_=9.2 MPa, E_L_=63 MPa) bringing it closer to results of SCW-T in all directions. This reinforces the geometrical pattern idealization and the NET approach as a suitable method to model the SCW.

##### Introduction of tortuosity into the NET model

The ~25-fold discrepancy in the longitudinal/axial moduli between NET and MAP led us to study the experimental map in more detail. We realized that microfibrils had a somewhat irregular, wavy appearance, with an uneven distribution of hemicellulose along the cellulose core, which results in a weakening of the microfibril in the axial direction. We modelled such imperfections by introducing a tortuosity (twist) in our idealized microfibril model. Introduction of tortuosity has a very limited influence in the transverse direction, but results in stiffness drop in the axial direction, which brings E_L_ in close agreement with the MAP model. We analyzed the implications of tortuosity by performing nonlinear buckling and viscoelastic analyses (Fig. 4H, I). While the buckling load is reduced ~50-fold, tortuous fibers are much more ductile as they can undergo ~9-times larger deformations before collapse in comparison to straight idealized fibers, storing up to 43% more elastic energy than straight fibers as they deform. We conclude that while tortuosity significantly reduces the stiffness of the cell wall, it also increases its ductility as well as the dissipation of elastic and viscoelastic energies, which could be crucial for plants to prevent breakage against extreme loading (such as in high winds).

##### Mechanical Analysis of NET model

We tested several geometrical features of the NET model, including network size, cross-link arrangement, number of cross-links with the same volume of hemicellulose, the inclination of the cross links as well the orientation of the microfibrils, and found that neither an increase of the height, nor length, nor thickness of the overall network significantly altered the stiffness or the relative elastic energy storage, if the ratio of cross links per length is kept constant, reinforcing the notion that mechanical analysis of relatively small cell wall portions is still relevant for entire cells. Fewer but thicker cross-links promote better load transfer to microfibrils, leading to more evenly distributed strain energy. The inclination of the hemicellulose cross-links had little effect on the overall stiffness.

## LITERATURE CITED

1. Van Iterson, G., 1937. A few observations on the hairs of the stamens of *Tradescantia virginica*. Protoplasma 27: 190–211.

2. Roelofsen, P.A. and Houwink, A. L. 1951. Cell wall structure of staminal hairs of *Tradescantia virginica* and its relation with growth. Protoplasma 40: 1–22.

3. Roelofsen, P. A., 1958. Cell-wall structure as related to surface growth. Some supplementary remarks multinet growth. Acta. Bot. Neerl. 7: 77–89.

4. Ohad I, Danon D (1964) On the dimensions of cellulose microfibrils. J Cell Biol 22: 302–305

5. Frey-Wyssling A (1968) The ultrastructure of wood. Wood Sci Technol 2: 73–83

6. Heyn AN (1969) The elementary fibril and supermolecular structure of cellulose in soft wood fiber. J Ultrast Res 26: 52–68

7. Somerville, C., Bauer, S., Brininstool, G., Facette, M., Hamann, T., Milne, J., Osborne, E., Paredez, A., Persson, S., Raab, T., Vorwerk, S. and Youngs, H. 2004. Toward a systems approach to understanding plant cell walls. Science 306: 2206–2211.

8. Cosgrove, D. J. 2005. Growth of the plant cell wall. Nature rev. Molecular cell biology 6: 850–861.,

9. Ding, S-Y. and Himmel, M. E. 2006. The maize primary cell wall microfibril: a new model derived from direct visualization. J. Agri. Food Chem. 54: 597–606.;

10. Ding SY, Liu YS, Zeng Y, Himmel ME, Baker JO, et al. (2012) How does plant cell wall nanoscale architecture correlate with enzymatic digestibility? Science 338: 1055–1060

11. McCann, M. C. and Roberts, K. 1991. Architecture of the primary cell wall. Academica Press, London, UK

12. Rose, J. K. C. and Bennett, A. B. 1999. Cooperative disassembly of the cellulose xyloglucan network of plant cell walls: parallels between cell expansion and fruit ripening. Trends Plant Sci. 4: 176–183.

13. Newman, R. H., Hill, S. J. and Harris, P. J. 2013. Wide-angle x-ray scattering and solid state nuclear magnetic resonance data combined to test models for cellulose microfibrils in mung bean cell walls. Plant Phys 163: 1558–1567

14. Thomas, L. H., Forsyth, V. T., Sturcová, A., Kennedy, C. J., May RP, Altaner, C. M., Apperley, D. C., Wess, T. J. and Jarvis, M. C. 2013. Structure of cellulose microfibrils in primary cell walls from collenchyma. Plant Physiol. 161: 465–476.

15. Sarkar, P., Bosneaga, E. and Auer, M. 2009. Plant cell walls throughout evolution: towards a molecular understanding of their design principles. Journal of Experimental Botany 60: 3615–3635.

16. Banasiak, A. 2014 Evolution of the cell wall components during terrestrialization Acta Soc Bot Pol 83(4):349–362

17. Llyod C. 2011. Dynamic microtubules and the texture of plant cell walls. Int. Rev. Cell Mol. Biol. 287: 287–329.

18. Sarkar, P. and Auer, M. (2016) Organization of the plant cell wall, In: Molecular cell biology of the growth and differentiation of plant cells, Rose, R. J [Ed.]: 101–119

19. Park, Y. B. and Cosgrove, D. J. 2012. A revised architecture of primary cell walls based on biomechanical changes induced by substrate-specific endoglucanases. Plant Physiol. 158: 1933–1943

20. Zienkiewicz, Olgierd Cecil, et al. The finite element method. Vol. 3. London: McGraw-hill, 1977.

21. Kha H, Tuble SC, Kalyanasundaram S, Williamson RE (2010) WallGen, software to construct layered cellulose-hemicellulose networks and predict their small deformation mechanics. Plant Physiol 152: 774–786

22. Yi, H., & Puri, V. M. (2012). Architecture-based multiscale computational modeling of plant cell wall mechanics to examine the hydrogen-bonding hypothesis of the cell wall network structure model. Plant physiology, 160(3), 1281–1292.

23. Nili, A., Yi, H., Crespi, V. H., & Puri, V. M. (2015). Examination of biological hotspot hypothesis of primary cell wall using a computational cell wall network model. Cellulose, 22(2), 1027–1038.

24. Sarkar, P., Bosneaga, E., Yap, E. G. Jr., Das, J., Tsai, W-T., Cabal, A., Neuhaus, E., Maji, D., Kumar, S., Joo, M., Yakovlev, S., Csencsits, R., Yu, Z., Bajaj, C., Downing, K. H. and Auer, M. 2014. Electron Tomography of Cryo-Immobilized Plant Tissue: a novel approach to studying 3d macromolecular architecture of mature plant cell walls in situ. PLoS ONE, 9: e106928.

25. The ultrastructural demonstration o/ compounds containing 1,2-glycol groups in plant cell walls G. G. JEWELL and c. A. SAXTON Histochemical Journal, 2 (I97O), 17–27

26. Zablackis E, Jing H, Muller B, Darvill AG, Albersheim P: Characterization of the cell-wall polysaccharides of Arabidopsis thaliana leaves. Plant Physiol 1995, 107: 1129–1138.

27. Busse-Wicher M, Gomes TC, Tryfona T, Nikolovski N, Stott K, Grantham NJ, Bolam DN, Skaf MS, Dupree P. The pattern of xylan acetylation suggests xylan may interact with cellulose microfibrils as a twofold helical screw in the secondary plant cell wall of Arabidopsis thaliana. Plant J. 2014; 79: 492–506

28. Simmons TJ, Mortimer JC, Bernadinelli OD, Poeppler AC, Brown SP, deAzevedo ER, Dupree R, Dupree P. Folding of xylan onto cellulose fibrils in plant cell walls revealed by solid-state NMR. Nature Communications 2016; 13902

29. Bailie, J. A., Ley, R. P., & Pasricha, A. (1997). A summary and review of composite laminate design guidelines, Northrop Grumman Report under NASA Contract NAS1-19347.

30. Park, Robert, and Thomas Paulay. Reinforced concrete structures. John Wiley & Sons, 1975.

31. Yan, Libo, and Nawawi Chouw. “Crashworthiness characteristics of flax fibre reinforced epoxy tubes for energy absorption application.” Materials & Design51 (2013): 629–640.

## Supplemental Material References

32. Yakovlev S, Downing KH (2011) Freezing in sealed capillaries for preparation of frozen hydratedsections. J Microscopy 244: 235–247

33. Suloway C, Pulokas J, Fellmann D, Cheng A, Guerra F, et al. (2005) Automated molecular microscopy: the new Leginon system. J Struct Biol 151: 41–60

34. Mastronarde DN (2005) Automated electron microscope tomography using robust prediction of specimen movements. J Struct Biol 152: 36–51

35. Kremer JR, Mastronarde DN, McIntosh JR (1996) Computer visualization of three-dimensional image data using IMOD. J Struct Biol 116: 71–76

36. Mastronarde DN (1997) Dual-axis tomography: an approach with alignment methods that preserve resolution. J Struct Biol 120: 343–352

37. Timoshenko, S. P., 1921, On the correction factor for shear of the differential equation for transverse vibrations of bars of uniform cross-section, Philosophical Magazine, p. 744.

38. Timoshenko, S. P., 1922, On the transverse vibrations of bars of uniform cross-section, Philosophical Magazine, p. 125.

